# STAG2 cohesin is essential for heart morphogenesis

**DOI:** 10.1101/788158

**Authors:** M. De Koninck, E. Lapi, C. Badia-Careaga, I. Cossio, D. Giménez-Llorente, M. Rodríguez-Corsino, E. Andrada, A. Hidalgo, M. Manzanares, F. X. Real, A. Losada

## Abstract

The distinct functions of cohesin complexes carrying STAG1 or STAG2 need to be unraveled. *STAG2* is commonly mutated in cancer and germline mutations have been identified in cohesinopathy patients. To better understand the underlying pathogenic mechanisms, we here report the consequence of *Stag2* ablation in mice. STAG2 is largely dispensable in adults and its tissue-wide inactivation does not lead to tumors but reduces fitness and affects both hematopoiesis and intestinal homeostasis. STAG2 is also dispensable for murine embryonic fibroblasts *in vitro*. In contrast, null embryos die by mid gestation showing global developmental delay and heart defects. Histopathological analysis and RNA-sequencing unveiled that STAG2 is required both for proliferation and regulation of cardiac transcriptional programs and in its absence, secondary heart field progenitors fail to enter the heart tube. These results provide compelling evidence on cell- and tissue-specific roles of the two cohesin complexes and how their dysfunction contributes to disease.

## Introduction

Cohesin is a four-subunit complex that holds the sister chromatids together to ensure faithful DNA repair by homologous recombination and proper chromosome segregation during cell division (*1*). It is present in all cells and its cohesive function is essential for proliferation. In addition, cohesin contributes to the spatial organization of the genome and to the activation and repression of tissue-specific transcriptional programs together with architectural proteins such as CTCF and transcriptional regulators like Mediator (*2*–*4*). In the cohesin complexes present in vertebrate somatic cells, the Structural Maintenance of Chromosomes (SMC) heterodimer of SMC1A and SMC3 associates with the kleisin subunit RAD21 and with one of the two versions of the Stromal Antigen (SA/STAG) subunit, STAG1 or STAG2. The two variants are present in all tissues and cell types, but their functional specificity is not well established (*5*). We previously showed that genetic ablation of *Stag1* in mice is embryonic lethal, which indicates that the two complexes are not redundant, at least during embryonic development (*6*). Lethality starts after day 11.5 of gestation (E11.5) but a small fraction of embryos survive to E18.5 and present a severe developmental delay and general hypoplasia (*7*).

In *Stag*1 null mouse embryonic fibroblasts (MEFs) telomere cohesion is impaired, preventing efficient replication of telomeres and causing chromosome segregation defects in mitosis (*6*). Centromere and arm cohesion are not clearly affected, which suggests that cohesin-STAG1 is specifically required for telomere cohesion whereas cohesin-STAG2 contributes to cohesion in other chromosomal regions. Results in human cells are in line with these findings, although the extent of cohesion defects reported in the absence of STAG2 is variable. In any case, a single variant is sufficient to maintain cell viability in culture (*6, 8, 9*).

Cohesin variants also contribute distinctly to genome organization and gene regulation. In *Stag1* null MEFs cohesin distribution is altered and so is their transcriptome (*7*). In the pancreas of heterozygous *Stag1* mice, the architecture of the *Reg* locus and the transcription of some of its genes are also changed compared to pancreas from wild type littermates, suggesting that STAG2 is not sufficient to compensate for the reduced levels of STAG1 (*10*). In human mammary epithelial cells, downregulation of STAG1 or STAG2 result in distinct changes in gene expression and chromatin contacts (*5*). Cohesin-STAG1 and cohesin-STAG2 colocalize with CTCF and play a major role in preservation of topologically associating domain (TAD) borders. By contrast, cohesin-STAG2 is also present at enhancers lacking CTCF that are bound by other transcriptional regulators (*3, 5, 11*). Importantly, cohesin-STAG1 cannot occupy these non-CTCF cohesin-sites even when STAG2 is absent (*5*). In mouse embryonic stem cells, cohesin-STAG2 promotes compaction of Polycomb domains and the establishment of long-range interaction networks between distant Polycomb-bound promoters that are important for gene repression (*11*).

Germline mutations in genes encoding cohesin and its regulatory factors are at the origin of a group of human syndromes collectively known as cohesinopathies. Cornelia de Lange syndrome (CdLS) is the most common of them and up to 60% of the patients carry heterozygous mutations in *NIPBL*, a protein involved in loading cohesin on chromatin (*12*). Clinical features often include growth retardation, intellectual disability, facial dysmorphism and congenital heart defects. Recently, germline mutations in *STAG1* and *STAG2* have been identified in patients with features partially overlapping those of CdLS and other cohesinopathies (*13*–*17*). Somatic mutations in cohesin genes, particularly in *STAG2*, have also been identified in several tumor types (*18*). *STAG2* has been recognized as one of the twelve genes significantly mutated in four or more cancer types (*19*). Among them, STAG2 loss is most frequent in urothelial bladder cancer (*20*). The evidence emerging from the study of diseases associated with both germline and somatic cohesin mutations strongly suggests that gene deregulation, rather than defects in chromosome segregation, underlies the pathogenic mechanism (*12, 20, 21*).

Given the growing importance of *STAG2* in human disease, we generated a *Stag2* conditional knock out (cKO) mouse strain to identify the consequences of eliminating cohesin-STAG2 in cells, embryos and adult tissues.

## Results

### Mild cohesion defects in STAG2 deficient MEFs

To generate a cKO allele of the X-linked *Stag2* gene, we used a targeting construct that carries loxP sites flanking exon 7 along with an FRT-flanked cassette encompassing a neomycin resistance selection gene, a splicing acceptor site and a polyadenylation sequence (Fig. S1A). Correctly targeted embryonic stem (ES) cells were screened by Southern blotting and were infected with adeno-FLP to eliminate this cassette before microinjection in C57BL/6J blastocysts (Fig. S1B). Germline transmission of the *Stag2*^lox^ allele was assessed by PCR (Fig. S1C). Next, *Stag2*^lox/lox^ females were crossed with males carrying *hUBC-CreERT2* for ubiquitous, tamoxifen-induced activation of the Cre recombinase. Embryos were extracted at embryonic stage E12.5 to generate cKO MEFs. Upon addition of 4-hydroxy-tamoxifen (4-OHT) to the culture medium for 4 days, STAG2 protein levels in treated MEFs (KO) dropped below 5% of the amount present in untreated MEFs (WT) and compensatory upregulation of STAG1 could be observed (Fig. 1A). The doubling time of STAG2 deficient cells was higher (Fig. 1B), but flow cytometry analysis did not reveal differences in the cell cycle profiles of WT and KO MEFs (Fig. 1C). We next examined cohesion and chromosome segregation. For these experiments, *Stag2* was deleted in serum-starved conditions and cells going through the first mitosis after release from the G0 arrest were collected. We detected very few cases of complete sister chromatid unpairing in metaphase spreads in WT or KO MEFs (1.3% and 3% of chromosomes examined, respectively; dark green square in Fig. 1D). However, we did observe a larger fraction of chromosomes per metaphase in which centromere cohesion was loosened, as evidenced by increased distance between sister centromeres (26% in KO vs 11% in WT MEFs; light green square in Fig. 1D). We also found a ca. 2-fold increase in the percentage of anaphase cells with lagging chromosomes and/or chromosome bridges among KO MEFs compared with WT MEFs (29% vs 17%), although the difference did not reach statistical significance (Fig. 1E). Finally, we observed that the proportion of metaphases with a normal chromosome number (n=40) decreased more the longer MEFs were grown in the absence of STAG2 (Fig. 1F). Overall, these defects are milder than those identified in *Stag1* null MEFs (*6*) or in C2C12 myoblasts or HeLa cells after STAG2 downregulation by siRNA (*6, 22*). We conclude that primary cultured cells almost completely lacking cohesin-STAG2 can proliferate, although at slower rates, and maintain sufficient cohesion to divide successfully but lose chromosomes more frequently that wild type cells.

**Fig. 1.**
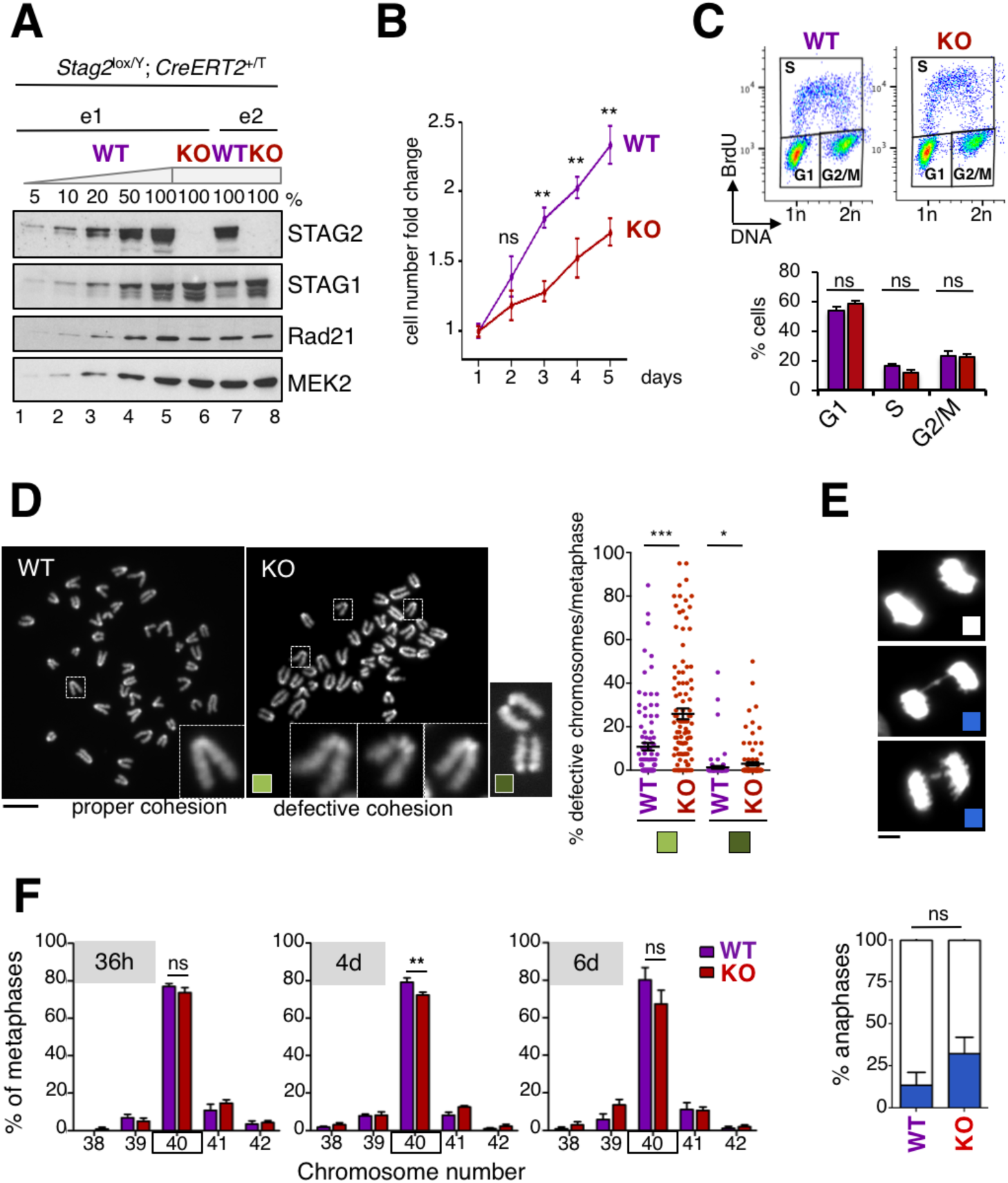
STAG2 deficient MEFs display slower proliferation and mild cohesion defects. **A.** Immunoblot analyses of whole cell extracts of *Stag2* cKO MEFs from two embryos (e1 and e2) untreated or treated with 4-OHT to activate CreERT2 (WT and KO hereafter). Decreasing amounts (shown as % of maximal) of WT MEF extract were loaded to estimate STAG2 depletion levels. MEK2 is used as loading control. **B.** Growth curves of WT and KO MEFs representing the average fold increase in cell number relative to the number of cells seeded on day 1. Data from MEFs from 2 embryos, each analyzed in triplicates (mean ± SEM). **C.** Representative BrdU incorporation profiles by FACS in WT and KO MEFs and bar graph showing values for n=4 (mean ± SEM). **D.** Representative metaphase spreads from WT and KO MEFs (left) and quantification (right, mean ± SEM) of centromeric cohesion defects. Each dot represents a single metaphase. At least 100 metaphases from MEFs from 3 different embryos were inspected. Scale bar, 10 µm. **E.** Images of anaphase cells, either normal or defective, found among WT and KO MEFs (top) and their quantification (bottom, mean ± SEM). Defective anaphases include lagging chromosomes and bridges. At least 100 anaphases from MEFs from 3 different embryos were inspected. Scale bar, 5 µm. **F.** Quantification of chromosome number frequency in metaphase spreads of WT and KO MEFs (mean ± SEM). The different timepoints (36h, 4d, 6d) refer to the time the cells have been cycling in the presence of 4-OHT; i.e. “36h” refers to 3 days of 4-OHT treatment upon serum starvation for proper STAG2 depletion, followed by 36h of release from the arrest. “4d” and “6d” indicate number of days in the presence of 4-OHT of asynchronously growing cells. At least 100 metaphases from MEFs from 3 different embryos were inspected. Mann-Whitney test; *** P<0.001, ** P<0.01, * P<0.05, ns P≥0.05.

### STAG2 is required for hematopoietic maintenance in adult mice

To determine whether STAG2 is essential in adulthood, 4-week old *Stag2* cKO mice carrying or not the *hUBC-CreERT2* transgene (KO and WT hereafter) were fed continuously with a tamoxifen-containing diet (TMX). We did not observe an acute loss of viability in the KO mice, but their survival was significantly shorter than that of WT mice (Fig. 2A). There were no preneoplastic or neoplastic lesions in the full necropsies of the mice analyzed (n=8). Likewise, a macroscopic assessment failed to reveal neoplasms in a large cohort of mice (n=63) of up to 70 weeks age, indicating that *Stag2* inactivation does not increase spontaneous tumor incidence in adult mice. At 12 weeks, loss of STAG2 protein was confirmed by immunohistochemistry in a large fraction of cells (>80%) of all tissues analyzed (Fig. 2B, left panel; Fig. 2C). However, over time, the fraction of STAG2-negative cells dropped dramatically in the more proliferative tissues (e.g. intestine and spleen) and to a lesser extent in tissues with lower proliferation rates such as liver, pancreas or brain (Fig. 2B and 2C). In the peripheral blood of KO animals carrying a dual fluorescent Cre reporter (*Rosa26_ACTB-tdTomato_EGFP*), the population of recombined leukocytes (GFP+, STAG2-) decreased with time (Fig. 2D).

**Fig. 2.**
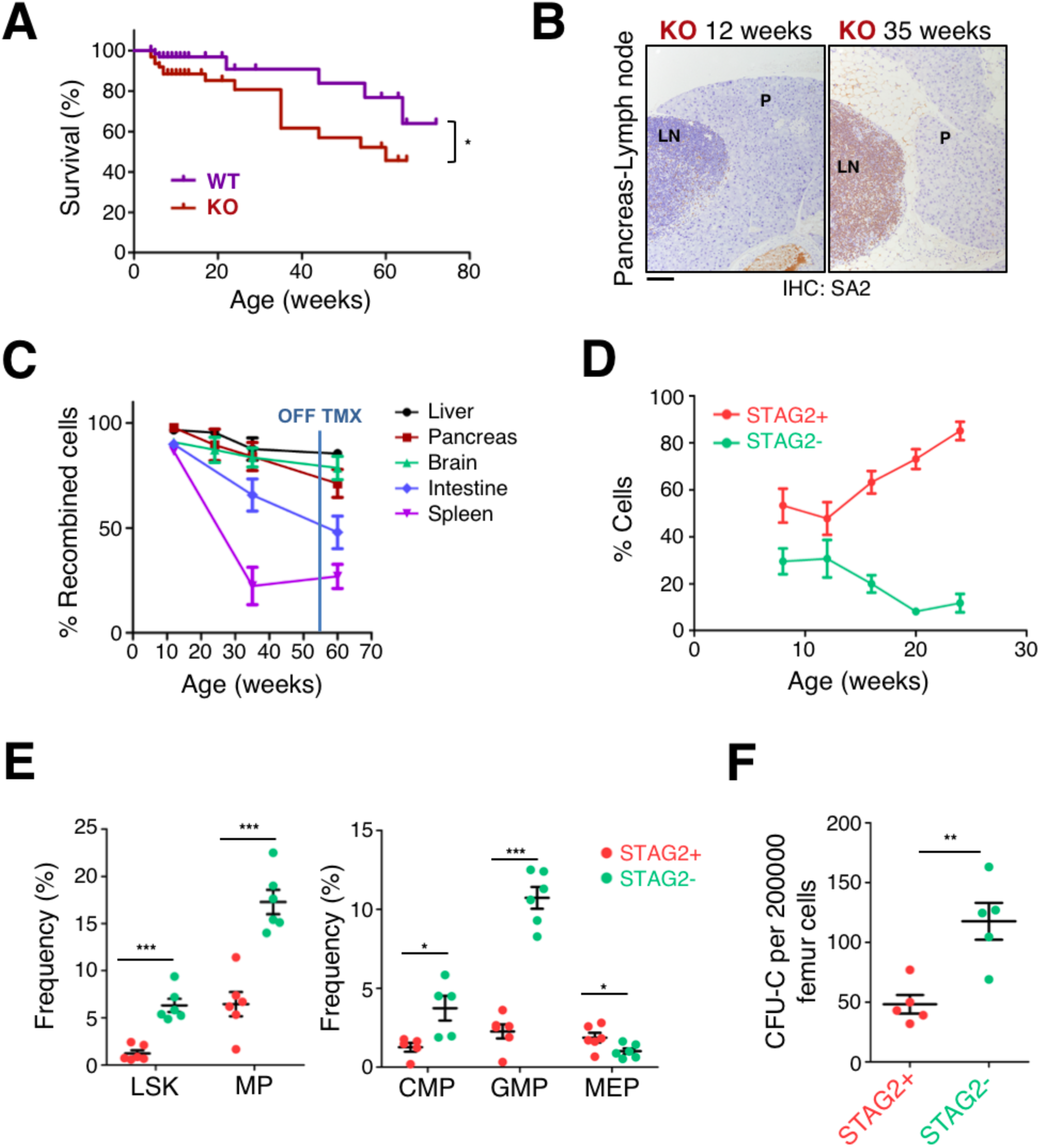
Effects of STAG2 ablation in adult mice. **A.** Kaplan-Meier survival curves of *Stag2* KO and WT mice. Four-week old *Stag2* cKO male and female mice carrying the *Cre-ERT2* allele (KO, n=63), or not (WT, n=66), were continuously fed on a TMX-containing diet and monitored thrice weekly. No gender differences in survival were observed. Gehan-Breslow-Wilcoxon test; *P < 0.05. **B.** Representative images of STAG2 expression in sections of pancreas (P) and associated lymph node (LN) of KO mice at 12 (left) and 35 (right) weeks of age, assessed by IHC. Scale bar, 100 µm. **C.** Percentage of recombined STAG2-negative cells in various organs over time was assessed by IHC. Representative microphotographs were quantified with Image J software. Error bars indicate SEM. **D.** Flow cytometry analysis of GFP+ (STAG2-) and Tomato+ (STAG2+) leukocytes in peripheral blood of KO mice over time (n=5). Error bars indicate SEM. **E.** Flow cytometry analysis of bone marrow HSPCs in 12 week-old KO mice (n=6). Left, LSK (Lin- c-Kit+ Sca1+); MP (Lin- c-Kit+). Right, CMP (Lin- c-Kit+ CD34+ CD1632-); GMP (Lin- c-Kit+ CD34+ CD1632+); MEP (Lin- c-Kit+ CD34- CD1632-). Error bars indicate SEM. Unpaired t test; *P < 0.05; ***P < 0.001 **F.** Colony-forming unit assay using FACS-sorted GFP and Tomato total bone marrow cells from 12 week-old KO mice (n=5). Error bars indicate SEM. Unpaired t test; **P < 0.01.

Upon treatment with TMX from weeks 4 to 12, KO mice displayed mild reductions in peripheral blood leukocyte, erythrocyte, and platelet counts pointing to anomalies in hematopoiesis (Fig. S2A,B). Analyses of leukocytes from peripheral blood and spleen of KO animals carrying the fluorescent Cre reporter revealed an enrichment in myeloid cells (monocytes and neutrophils) and a significant reduction in T lymphocytes among STAG2 deficient cells compared with unrecombined cells (Fig. S2C). In bone marrow, a clear expansion of the LSK (Lin-Sca1+ c-Kit+) and myeloid progenitor (MP) compartment was associated with STAG2 loss (Fig. 2E, left). Further analysis of committed progenitors revealed increased frequency in CMPs (common myeloid progenitor) and GMPs (granulocyte-monocyte progenitor) and a decrease in MEPs (megakaryocyte-erythrocyte progenitor) among STAG2 deficient cells, in agreement with the findings in peripheral blood (Fig. 2E, right). Reductions in MEP paralleled a decrease in bone marrow Ter119+ cells in KO mice (Fig. S2D,E). Functional analyses revealed a higher colony forming capacity of STAG2 deficient hematopoietic cells (Fig. 2F), while the loss in lymphoid potential explained the reduced chimerism of mutant cells over time in peripheral blood (Fig. 2D). These results support the notion that STAG2 loss leads to increased self-renewal and impaired differentiation of hematopoietic stem cells (HSC), myeloid skewing and an overall competitive disadvantage when wild type HSC are present. These data are also consistent with previous reports on the contribution of cohesin to normal hematopoiesis and the occurrence of cohesin mutations in myeloid malignancies (*21, 23, 24*).

### STAG2 is required for intestinal homeostasis

Shortly after the initiation of TMX diet at 4 weeks, the survival curve of KO mice diverted from that of WT mice, the former showing reduced body weight (Fig. 2A and Fig. 3A). Histological analyses at 8 weeks failed to reveal major alterations in tissues of KO mice with the exception of the intestine where patches of epithelial erosion and necrosis were observed, with moderate/severe lesions present in 60% of mice. Wild type mice showed intestinal lesions although they were much milder (Fig. 3B). We analyzed proliferation and apoptosis in the small bowel: intestinal crypts from KO mice showed a significant reduction of BrdU+ cells (Fig. 3C) and a significant increase of active caspase-3 labelling (Fig. 3D), suggesting reduced regeneration capacity. To acquire further insight into the requirement of STAG2 for intestinal cell renewal, we generated primary organoid cultures from the small intestine of TMX-treated KO mice carrying the fluorescent reporter (Figure 3E, left). GFP+ STAG2 null cells yielded fewer and smaller organoids than Tomato+ STAG2 proficient cells (Fig. 3E, right). These findings support the notion that STAG2 is required for intestinal homeostasis.

**Fig. 3.**
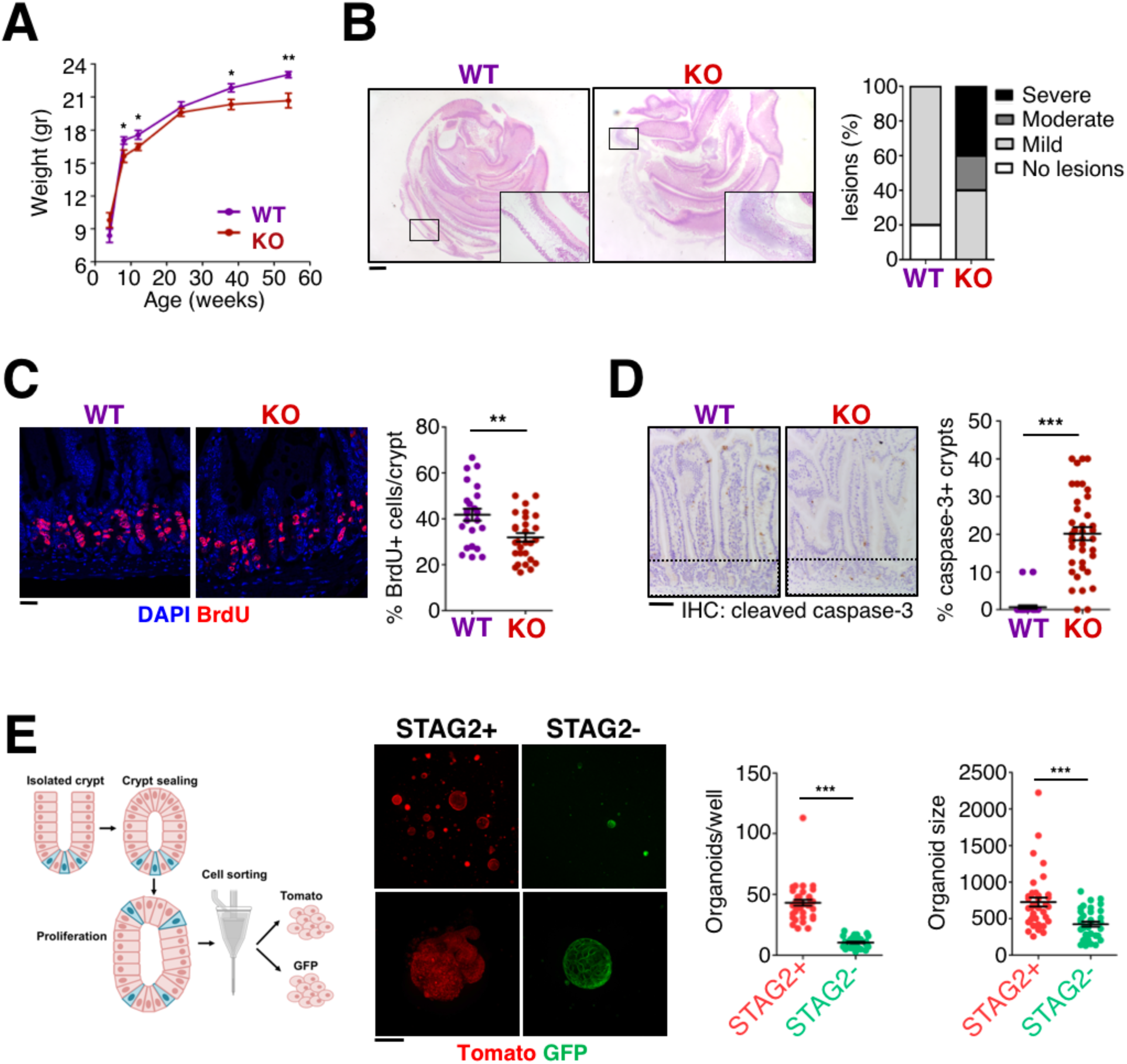
Requirement of STAG2 for intestinal cell renewal. **A.** Weight of KO and WT mice over time (n= 18 WT, n=18 KO). Error bars indicate SEM. One-sided Mann–Whitney U test; * P<0.05; **P < 0.01. **B.** Representative images of H-E-stained small intestine sections of 8 week-old WT and KO mice (left) and semi-quantitative assessment of severity of lesions (right). Scale bar, 1mm. (n= 5 WT and n=5 KO). **C.** Immunofluorescence analysis of BrdU (red) in sections of 8 week-old WT and KO intestine. Nuclei are counterstained with DAPI (blue). Scale bar, 25 µm. The percentage of BrdU+ cells per crypt is shown in the graph on the right (n= 24 WT, n=29 KO). Error bars indicate SEM. Two-tailed Mann–Whitney U test; **P < 0.01. **D.** Immunohistochemical analysis of cleaved caspase-3 in sections of 8 week-old WT and KO intestine. Nuclei are counterstained with hematoxylin. Crypt region is indicated by a dashed box. Scale bar, 50 µm. The percentage of crypts per section showing cleaved caspase-3 staining is plotted in the graph on the right (n= 30 WT, n=39 KO). Error bars indicate SEM. Two-tailed Mann–Whitney U test; ***P < 0.001. **E.** Scheme depicting the experimental design of intestinal organoid generation (left). Confocal microscopy images of Tomato+ (STAG2+) and GFP+ (STAG2-) organoids (middle). Scale bar, 100 µm. Quantification of the number and size of organoids (in pixels) obtained from cells sorted from primary intestinal organoids (5000 cells/well) (right). Error bars indicate SEM. Paired T test; ***P < 0.001.

### Stag2 null embryos display developmental delay by E9.5 and die soon afterwards

To determine whether STAG2 is essential for embryonic development, we crossed *Stag2*^lox/lox^ females with males carrying the *CAG-Cre* transgene, coding for a Cre recombinase that is expressed ubiquitously from the zygote stage. Since *Stag2* is an X-linked gene, male embryos resulting from this cross would be either wild type (WT) or null (KO) for *Stag2* while females would be WT or heterozygous (see genotyping strategy in Fig. S1D). We thus focused our analyses on the male embryos. None of the KO male embryos extracted at E12.5 was alive, suggesting an earlier embryonic lethality (Fig. 4A). In litters extracted at earlier stages we found live KO embryos at (or close to) the expected Mendelian ratios at E8.5 and E9.5, but not later (Fig. 4A). Immunohistochemical analyses of embryo sections with STAG2-specific antibodies confirmed tissue-wide absence of the protein (a section from the E9.5 embryonic heart is shown as an example in Fig. 4B). Importantly, E9.5 KO embryos were visibly smaller than their wild type littermates with variable penetrance of the phenotype (mild and severe examples shown in Fig. 4C). Despite no overt gross morphological defects, at least in embryos displaying the more prevalent milder phenotype, a significantly reduced number of somites was observed in mutant embryos starting at E9.5, indicating a clear developmental delay. By 10.5 this difference was equivalent to a 1-day lag in somite counts.(Fig. 4D). Thus, loss of STAG2 causes a generalized developmental delay, noticeable by E9.5 with variable penetrance, and results in death by E10.5.

**Fig. 4.**
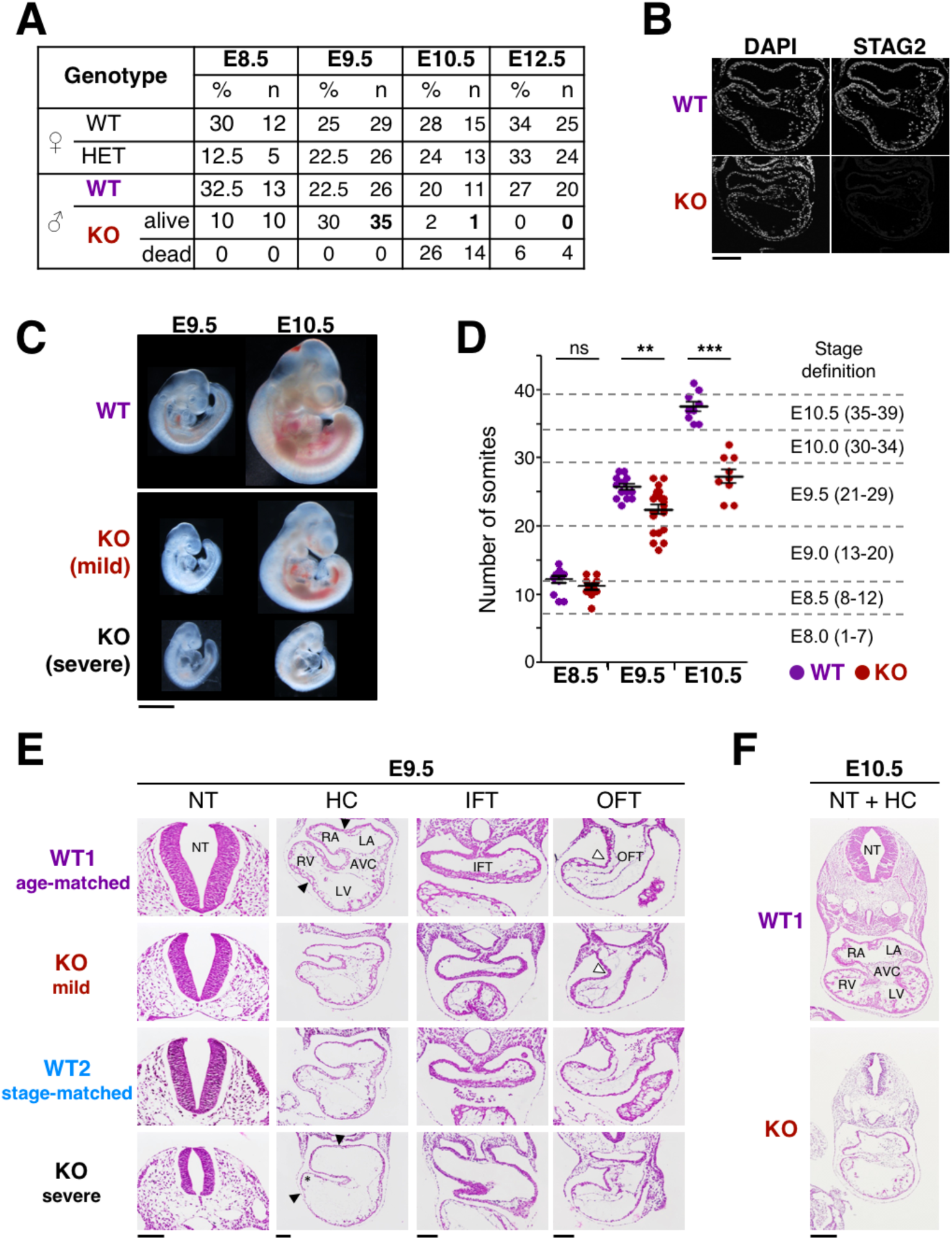
STAG2 becomes essential by mid-gestation and mutant embryos display heart defects. **A.** Viability of STAG2 deficient embryos at different stages of development. We extracted 6, 14, 7 and 13 litters at E8.5, E9.5, E10.5 and E12.5, respectively. Genotypes for *Stag2* are: female WT (lox/+), female HET (D/+), male WT (lox/Y), male KO (D/Y). **B.** Immunofluorescence staining of STAG2 in transverse heart sections of WT and KO embryos at E9.5. Nuclei are counterstained with DAPI. Scale bar, 200 µm. **C.** Representative images of WT and KO embryos (mild and severe phenotypes) at E9.5 and E10.5. Scale bar, 1mm. **D.** Somite number of WT and KO embryos at E8.5, E9.5 and E10.5. Somites were counted for embryos from 6 litters at E8.5 (n=13 WT and n=10 KO), 9 litters at E9.5 (n= 19 WT and n=25 KO) and 4 litters at E10.5 (n=10 WT and n=10 KO). Two-tailed Student’s t-test, *** P<0.001, ** P<0.01, ns P≥0.05. **E.** H-E stained transverse sections of KO (mild and severe), WT1 (age-matched control) and WT2 (stage-matched control) embryos extracted at E9.5; neural tube (NT), heart chambers (HC), inflow tract (IFT) and outflow tract (OFT); RA: right atrium; LA: left atrium; AVC: atrioventricular canal; RV: right ventricle; LV: left ventricle. Black arrowheads indicate the position of the prospective septum between right and left chambers. White arrowheads point at the OFT curve. Asterisk highlight the small size of the RV. Scale bars (valid for entire column), 100 µm. **F.** H-E stained transverse sections encompassing the neural tube and the heart of WT and KO embryos at E10.5. Heart regions indicated in WT, as in **E**. Scale bar, 250 µm.

### Aberrant heart morphogenesis in Stag2 null embryos

To identify developmental defects that could explain embryonic lethality, we analyzed the histology of E9.5 KO embryos with both severe and mild growth phenotypes and compared them with two different types of wild type controls: littermates (age-matched, WT1) and embryos from different litters but with the same number of somites (stage-matched, WT2). Mutants with a more severe phenotype showed aberrant morphology of several structures (neural tube, Fig. 4E; aorta, branchial arches, brain, otic and optic vesicles, Fig. S3). In contrast, most tissues and organs from KO embryos with a milder phenotype did not show obvious malformations but were clearly more similar to stage-matched than to age-matched controls (Fig. 4E; Fig. S3). A remarkable exception to this general trend was a selective defect in the developing heart: the prospective atria and right ventricle were reduced in size compared to both controls, while no clear differences were found in the left ventricle (compare images for WT1, KO mild and WT2 under HC, heart chambers, in Fig. 4E). Morphological defects were also observed in the outflow tract (OFT), where KO embryos showed an aberrant rightwards turning at the junction with the ventricular myocardium when compared to stage-matched controls (white arrowhead in WT1 and KO mild under OFT in Fig. 4E). However, the inflow tract (IFT) appeared normal (Fig. 4E). The defects described above were exacerbated in mutants with a severe phenotype, which displayed distended atria and ventricles with no visible indication of a future septum between right and left chambers (black arrowheads in WT1 and KO severe, under HC, in Fig. 4E), and abnormal right ventricle development (asterisk in Fig. 4E). In this case, both the OFT and the IFT were distended. Despite the variable penetrance of the phenotype by E9.5, all mutants displayed severe cardiac anomalies by E10.5, resembling those identified in the severe KO phenotype by E9.5, as well as extensive necrosis and apoptosis (Fig. 4F). Thus, defective heart function may account for the embryonic lethality of *Stag2* KO embryos.

### Decreased proliferation in Stag2 mutant embryos

To shed light into the cellular mechanisms leading to the defects described above, we first confirmed that both STAG1 and STAG2 are expressed in the heart of E9.5 wild type embryos using immunofluorescence (Fig. S4). These findings are consistent with data from single cell RNA sequencing of E8.25 murine embryos, which shows similar patterns of expression for both genes (*25*). We next analyzed proliferation and apoptosis in E9.5 WT1, WT2, and KO embryos with milder phenotype to uncover primary defects. Heart sections, as well as sections containing the neural tube for comparison, were labeled with anti-phosphohistone H3 (H3P) to detect proliferating cells and with Terminal deoxynucleotidyl transferase dUTP nick end labeling (TUNEL) to mark apoptotic cells. To identify the different heart compartments, co-labeling with Islet 1 (ISL1) was used: ISL1 is a transcription factor expressed throughout the anterior and posterior secondary heart field progenitors (ASHF and PSHF, respectively) and is progressively turned off in their descendants as they migrate into, and populate, the heart tube through its anterior and posterior ends (OFT and IFT, respectively, see scheme in Fig. S5A)(*26*). The fraction of H3P-positive cells in the heart chambers (atria and ventricles) was significantly lower in the mutant as compared to their littermate age-matched WT1 controls, but similar to WT2 stage-matched controls (HC in Fig. 5A, 5B, and Fig. S5B). These differences were reproduced in ASHF and OFT, as well as the neural tube, while they were less prominent in PSHF and IFT (Fig. 5A and 5B, Fig. S5B). There was also increased apoptosis in mutant neural tube and heart chambers as compared to both controls, although the number of TUNEL positive cells was very low in all cases and inter-individual variability high (Fig. S6). These results suggest that the global developmental delay observed in *Stag2* mutants at E9.5 is mainly due to a decrease in proliferative capacity of mutant cells.

**Fig. 5.**
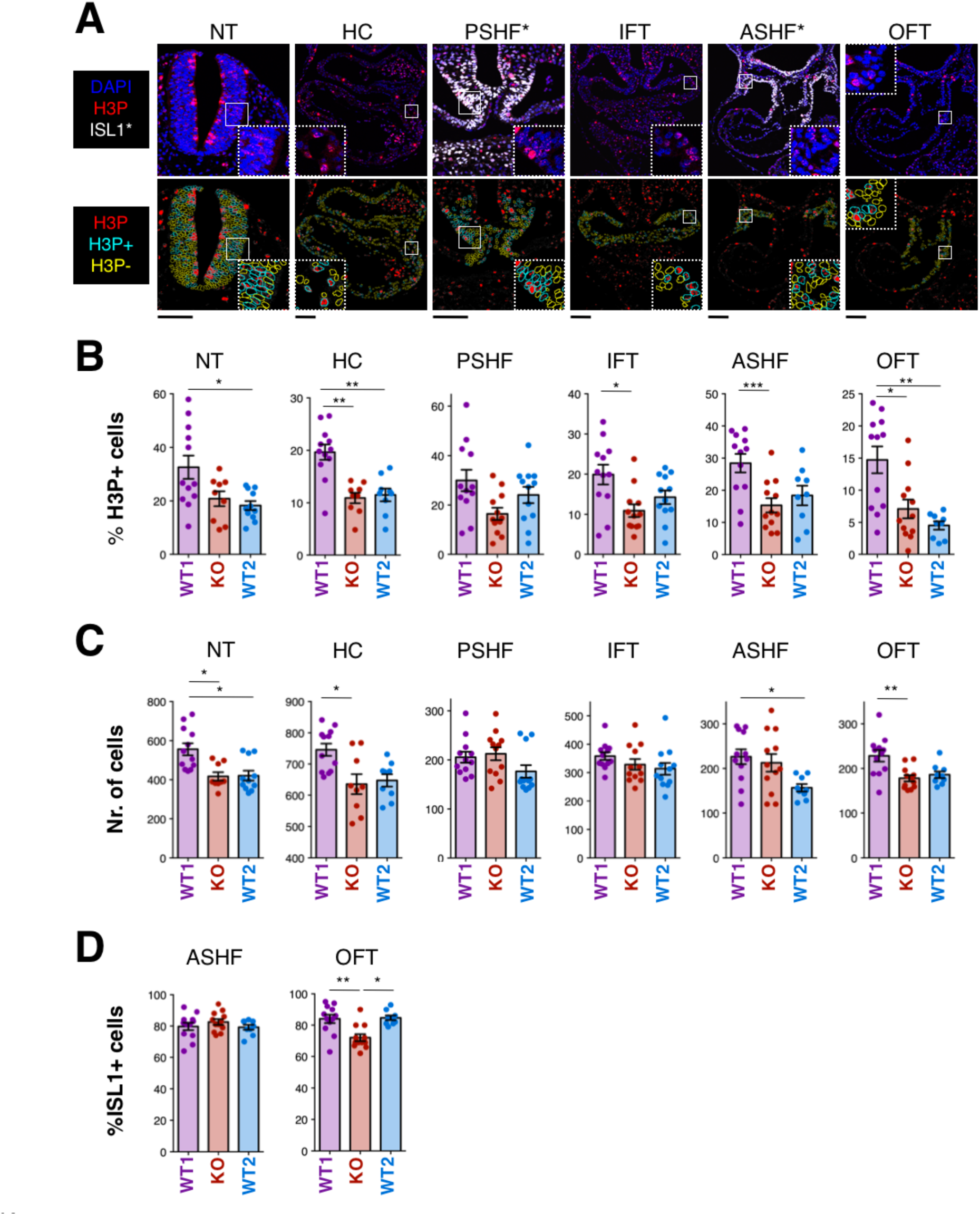
Reduced cell proliferation and impaired migration of ASHF progenitors in the developing heart of *Stag2* null embryos. **A.** Representative transverse sections of H3P staining of E9.5 WT1 embryos showing neural tube (NT), heart chambers (HC), posterior secondary heart field (PSHF), inflow tract (IFT), anterior secondary heart field (ASHF) and outflow tract (OFT). Upper row: original immunofluorescence signal of DAPI (blue), H3P (red) and ISL1 (white, only shown in regions marked with *). Lower row: H3P signal converted to a binary image, with representation of nuclei selection. Similar images for KO and WT2 embryos are shown in Fig. S5. Scale bars, 100 µm. **B.** Quantification of H3P-positive cells as readout for proliferation in E9.5 WT1 (age-matched control), KO (mild phenotype) and WT2 (stage-matched control) embryos in the indicated regions. 9-12 non-consecutive sections from 3-4 embryos were analyzed per genotype and region. **C.** Quantification of total number of cells per section in E9.5 WT1, KO and WT2 embryos in the indicated regions. **D.** Quantification of ISL1-positive cells in the ASHF and OFT regions of E9.5 WT1, KO and WT2 embryos.

### Impaired deployment of progenitors into the heart tube in Stag2 mutant embryos

While decreased proliferation might account for the global growth delay observed in the heart (and other organs) in mutant embryos, it failed to explain why morphological defects were more evident in certain heart structures, i.e. the OFT and right ventricle (Fig. 4E). Interestingly, these structures derive from second heart field (SHF) progenitors, while the left ventricle derives from the first heart field (FHF) (*27*). More specifically, ISL1+ progenitors present in the ASHF migrate into the heart tube contributing to the OFT and right ventricle (Fig. S5A). In mutant embryos, the length of inner and outer curves of the OFT was reduced compared to controls, suggesting a problem in the migration of ASHF progenitors (ISL1+) into the OFT (Fig. S7). To test this possibility, we quantified total cell numbers in heart sections. In the neural tube, heart chambers and OFT, cellularity of KO embryos was lower than in WT1 and more similar to WT2 embryos, consistent with reduced size and proliferation rates. In contrast, cell numbers in the ASHF were similar in KO and WT1 littermates (Fig. 5C), despite mutants showing a much reduced proliferation rate (Fig. 5B). Moreover, while the fraction of ISL1+ progenitors in ASHF was similar in all embryos, it decreased in the OFT of KO embryos compared to both controls (Fig. 5D). These findings strongly suggest that STAG2 loss results in accumulation of progenitors in the ASHF that fail to migrate into the heart tube, leading to morphological defects in ASHF derivatives such as the right ventricle and the OFT. Defects in migration of progenitors has been suggested as the cause of heart defects in murine embryos and zebrafish deficient for the cohesin loader NIPBL (*28, 29*).

### Altered transcription of cardiac development regulators in Stag2 mutant embryos

To address if the role of cohesin in gene regulation could contribute to the phenotypes described above, we compared the heart transcriptomes of E9.5 *Stag2* KO and WT embryos by RNA-seq. To exclude variation related to developmental stage, we selected littermate embryos of both genotypes with similar number of somites. To identify tissue-specific changes, we extracted RNA from the heart and from the neural tube lying adjacent to the heart. There were 1,881 differentially expressed genes (DEGs, FDR<0.05) between wild type samples of the two tissues, which defined a cardiac-enriched and a neural-enriched gene set (1,116 and 765 genes, respectively). Gene Ontology analysis confirmed the functional specificity of these gene sets (“cardiac” and “neural” genes, for simplicity; Fig. 6A and Table S1). STAG2 loss had a greater impact on the heart transcriptome, as shown in the heatmap of Fig. 6A. Accordingly, pairwise comparisons between WT and KO samples for each tissue identified 846 DEGs in heart but only 5 in neural tube (FDR<0.05; Fig. 6B and Table S2). Among the DEGs in heart, there were 222 and 112 genes from the cardiac and neural gene sets, respectively, indicating that tissue-specific genes were preferentially affected by STAG2 loss (Fig. 6C and Table S1). Moreover, most cardiac genes were downregulated in the heart of mutant embryos, whereas the neural genes were upregulated therein (Fig. 6D). These findings agree with the proposed role of cohesin-STAG2 in tissue-specific transcription, promoting activation of genes specifying a tissue (i.e., cardiac genes in heart) and repression of alternative gene programs (e.g., neural genes in heart) (*5*). A closer look at the list of DEGs in heart revealed several cardiomyocyte markers and well-established regulators of SHF among the downregulated genes (Fig. 6B, right). For instance, Fgf8 and Hand2 contribute to the survival of ASHF progenitors while Wnt5a activity is critical for their deployment into the OFT (*30*–*32*), consistent with the defects described in the previous section. Taken together, our data suggest an important role of STAG2 in the control of the early cardiac transcriptional programs that is not assumed by STAG1 in the *Stag2* KO embryos and which, together with decreased proliferation, contributes to the observed defects in heart morphogenesis.

**Fig. 6.**
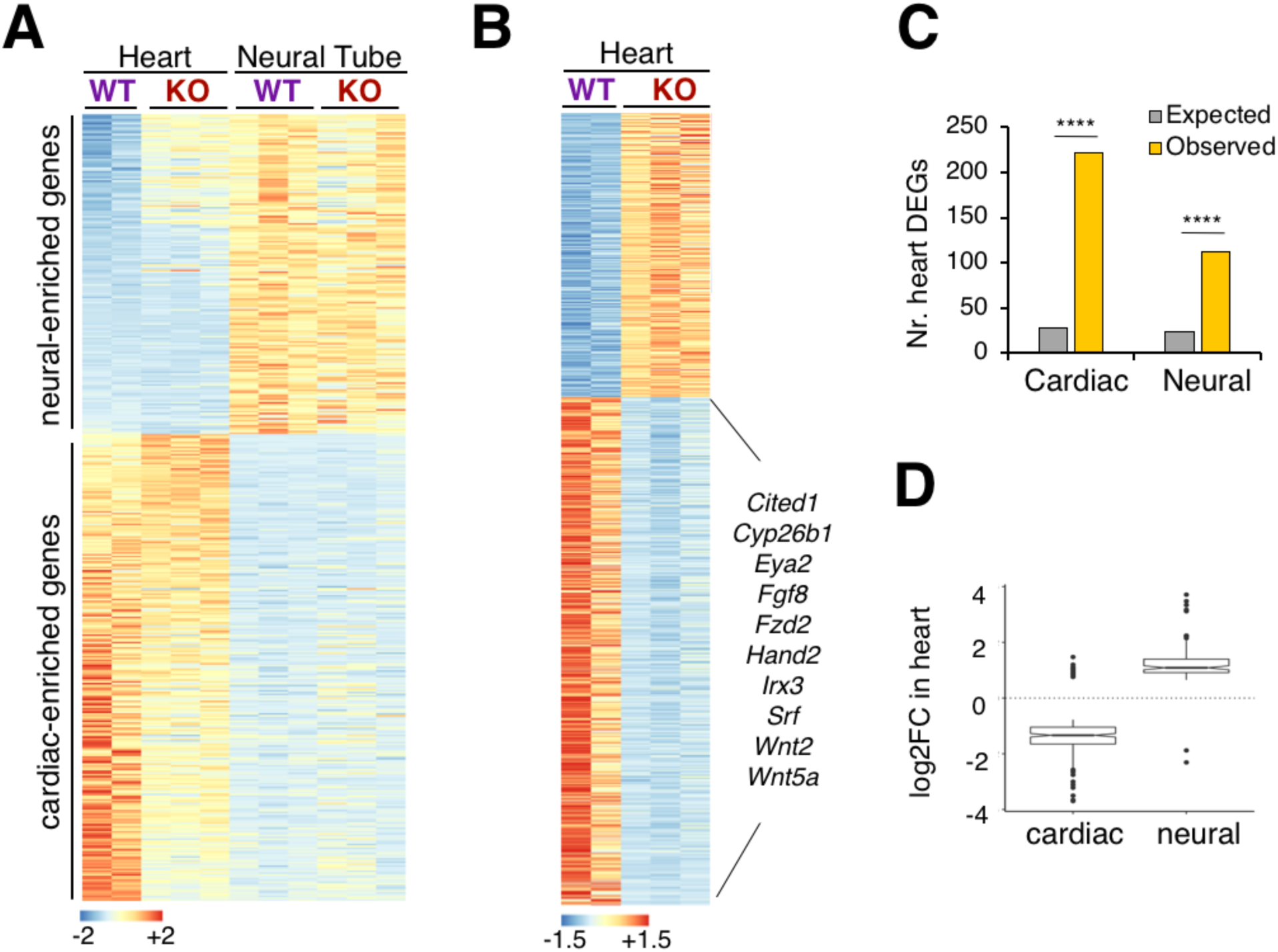
Transcriptional deregulation in *Stag2* null embryos. **A.** Heatmap showing relative expression of 1,116 cardiac- and 765 neural-enriched genes in all samples. Gene sets were defined by differential expression between WT heart and WT neural tube samples. **B.** Heatmap of 846 DEGs in WT and KO heart samples. Among the downregulated genes, we highlight some with established roles in cardiomyocyte differentiation and SHF. **C.** Expected versus observed number of cardiac and neural genes found among the heart DEGs. Total number of expressed genes: 21,653. Fisher’s exact test; ****<0.0001 (p<2E-12). **D.** Box plot of expression changes in the cardiac and neural genes identified as DEGs in heart. See also Table S1.

## Discussion

A major challenge in cohesin biology is to understand the specific functions of STAG1 and STAG2 (*5, 11*). To address this question, we previously characterized a *Stag1* KO mouse (*6, 7*) and we have now generated a *Stag2* KO mouse. While our manuscript was in revision, a study describing the consequences of *Stag2* ablation in the hematopoietic system was published, demonstrating a specific role for STAG2 in balancing self-renewal and differentiation in hematopoietic precursors (*24*). Here, we describe instead the consequences of ubiquitous STAG2 elimination in embryos and adult mice.

Whole body deletion of *Stag2* in young mice does not result in acute loss of viability, which suggests that STAG1 can largely compensate for the lack of STAG2 postnatally. Efficiency of Cre-mediated recombination of the *Stag2* cKO allele was high in adult tissues 8 weeks after Cre induction but the fraction of recombined cells decreased notably over the subsequent weeks in proliferative tissues despite continuous TMX administration. This observation indicates a clear proliferative disadvantage of STAG2 deficient cells and does not allow us to rule out that a more severe phenotype might be disclosed upon achieving a more complete or sustained depletion of STAG2. Consistent with results obtained upon *Stag2* deletion in HSC using *Mx1-Cre*, ubiquitous deletion also results in increased self-renewal and defective lineage commitment (*24*). Histopathological analyses also revealed defects in gastrointestinal tract homeostasis, most likely related to the toxicity of prolonged TMX administration from an early age (*33*) and the impaired regeneration capacity of *Stag2* KO crypt cells. Reduced proliferation was also found in *Stag2* null embryos, and probably contributes to their growth retardation. Our in vitro studies using *Stag2* null MEFs show loosened centromere cohesion that could lead to an increased rate of chromosome segregation defects and the generation of aneuploid, inviable cells thus contributing to reduced proliferation in vitro and in vivo.

In contrast to the redundancy and functional compensation of the STAG proteins in adult mice, embryos require both proteins to complete development. Constitutive inactivation of *Stag1* in the germline is embryonic lethal and causes severe development delay, with incomplete penetrance, but no obvious organ malformation (*6, 7*). In contrast, inactivation of *Stag2* in the germline leads to earlier lethality, starting at E9.5. This phenotype is associated with a broad, subtle, general tissue disorganization and a dramatic effect on heart development with no *Stag2* null embryos surviving beyond E10.5. It is unlikely that reduced proliferation alone accounts for these defects. Other mouse models partially deficient for genes important for cell proliferation, such as those carrying hypomorph alleles of the MCM replicative helicase, survive to later stages of embryo development and the associated lethality seemingly results from impaired expansion of hematopoietic precursors (*34*). Mutant mice in the centrosome component Cep57 that display more severe chromosome segregation anomalies than those reported here also survive to birth (*35*).

Increasing evidence supports the notion that the presence of cohesin-STAG2 at enhancer elements independently of CTCF promotes cell type-specific transcription, a function that is not compensated by cohesin-STAG1 (*5, 11, 24*). Consistent with this idea, our transcriptome analyses uncovered altered tissue-specific transcription patterns in *Stag2* null embryonic hearts, with lower expression of cardiac genes and de-repression of genes from other lineages. Thus, we propose that defects in both proliferation and lineage specification contribute to the heart abnormalities observed in the STAG2 deficient embryos, as previously suggested in NIPBL deficient embryos or zebrafish with reduced cohesin levels (*29, 36*).

Our results support a causative contribution of cohesin-STAG2 function to the cardiac anomalies detected in CdLS patients, most of them carrying *NIPBL* mutations (*37*). Cohesinopathy cases with *STAG2* mutations have been reported recently. Among those, male patients carry missense variants and show milder phenotypes that do not include heart defects while ventricular septal defects and other heart anomalies have been described in female patients carrying loss of function or missense variants (*13*–*17*). Since *STAG2* is an X-linked gene, the embryonic lethality of *Stag2* null murine embryos reported here explains why inactivating germline mutations will not be tolerated in males while heterozygous females may survive through the selection of cells in which the wild type allele is not silenced by the X inactivation process. The variable penetrance observed in STAG2 deficient mice as well as the contribution of epistatic events thus likely accounts for phenotypic diversity in patients carrying *STAG2* mutations.

In summary, we here show that cells lacking cohesin-STAG2 are viable both in vitro (tissue culture) and in vivo (in embryos and adult tissues), confirming that cohesin-STAG1 is sufficient to fulfill essential cohesin functions (*8, 9*). However, their decreased proliferation and altered transcriptomes lead to embryonic lethality, a result that provides further compelling evidence for cell- and tissue-specific roles of the two cohesin complexes and how their dysfunction may contribute to disease. We speculate that genomic changes derived from decreased accuracy of chromosome segregation and/or DNA repair as well as transcriptional alterations affecting cell identity may underlie the behavior of *STAG2* mutant tumors. While inactivation of *Stag2* in adult mice did not increase tumor incidence in our study, similar to other major tumor suppressor genes such as *Cdkn2a* or *Rb* (*38, 39*), these mice will be useful to model the cooperation of *STAG2* mutations with other genetic alterations in promoting tumorigenesis in a wide variety of cell types.

## Acknowledgments

We acknowledge the excellent technical support of the CNIO Transgenic Mouse Models unit led by Sagrario Ortega and the CNIC Microscopy unit, in particular Verónica Labrador, as well as the help of Natalia del Pozo, Ana Cuadrado, Alba de Martino, Eduardo Caleiras, and Cristian Perna.

## Author contributions

E.L. and M.R.C. generated the *Stag2* cKO mouse; M.D.K. characterized MEFs; E.L., E.A. and I.C. carried out the studies in adult mice; M.D.K. and C.B.C. performed the embryo studies; D.G.-LL. analyzed the RNA-seq data; A.H., M.M., F.X.R., and A.L. supervised the study and contributed to experimental design and data interpretation.

## Data and materials availability

Requests of *Stag2* cKO mice should be addressed to A. Losada and F. X. Real.

---

**Fig. S1.**
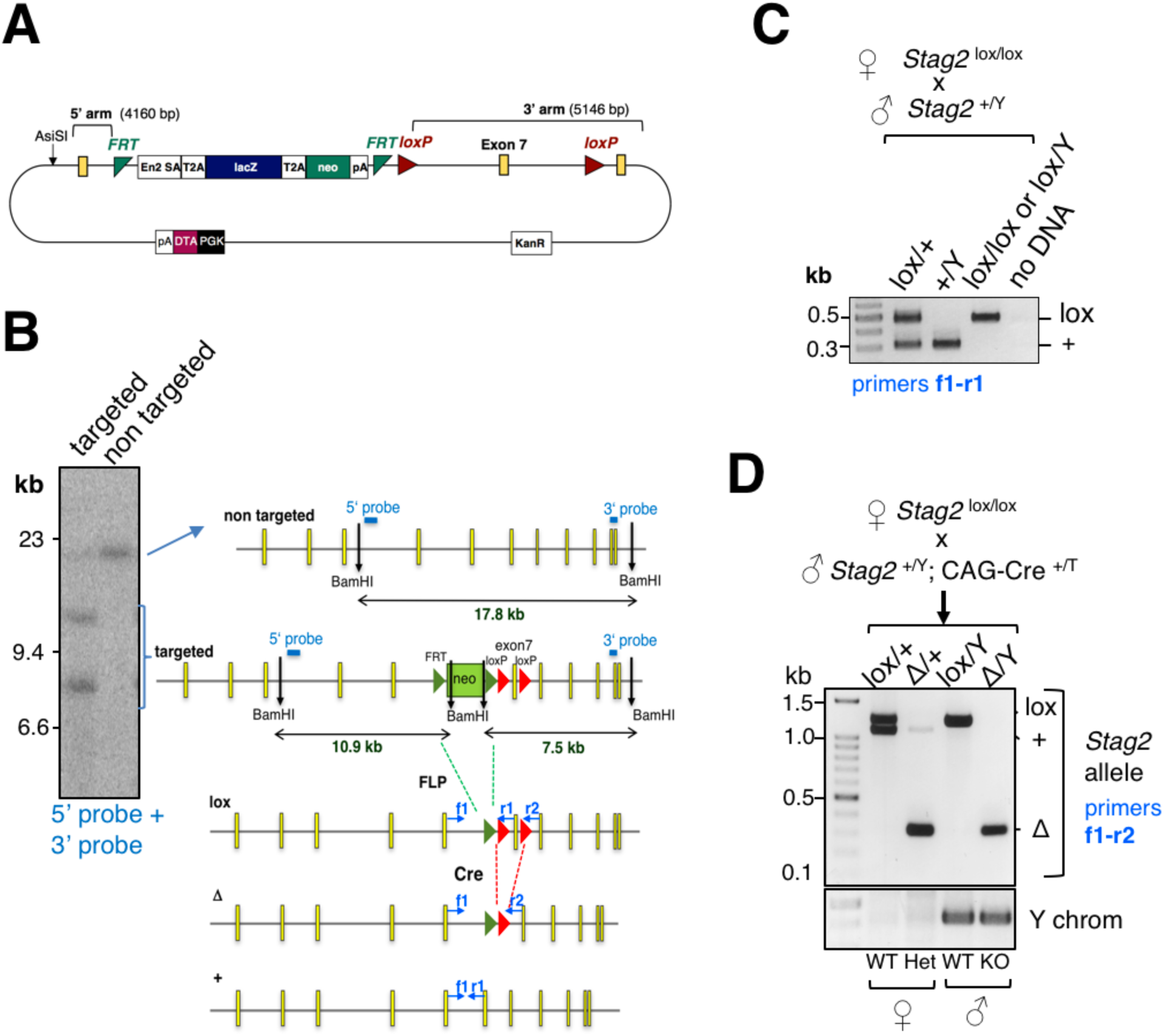
Generation of a *Stag2* cKO allele. **A.** Map of the vector obtained from EUCOMM to target the murine *Stag2* gene. **B.** Southern blot analysis (left) and strategy (right) to identify targeted ES clones. **C.** PCR analyses to genotype the *Stag2* lox and wild type (+) alleles in the offspring of the indicated mating. **D.** PCR analyses to genotype the embryos obtained from the indicated cross, including the Y chromosome.

**Figure S2.**
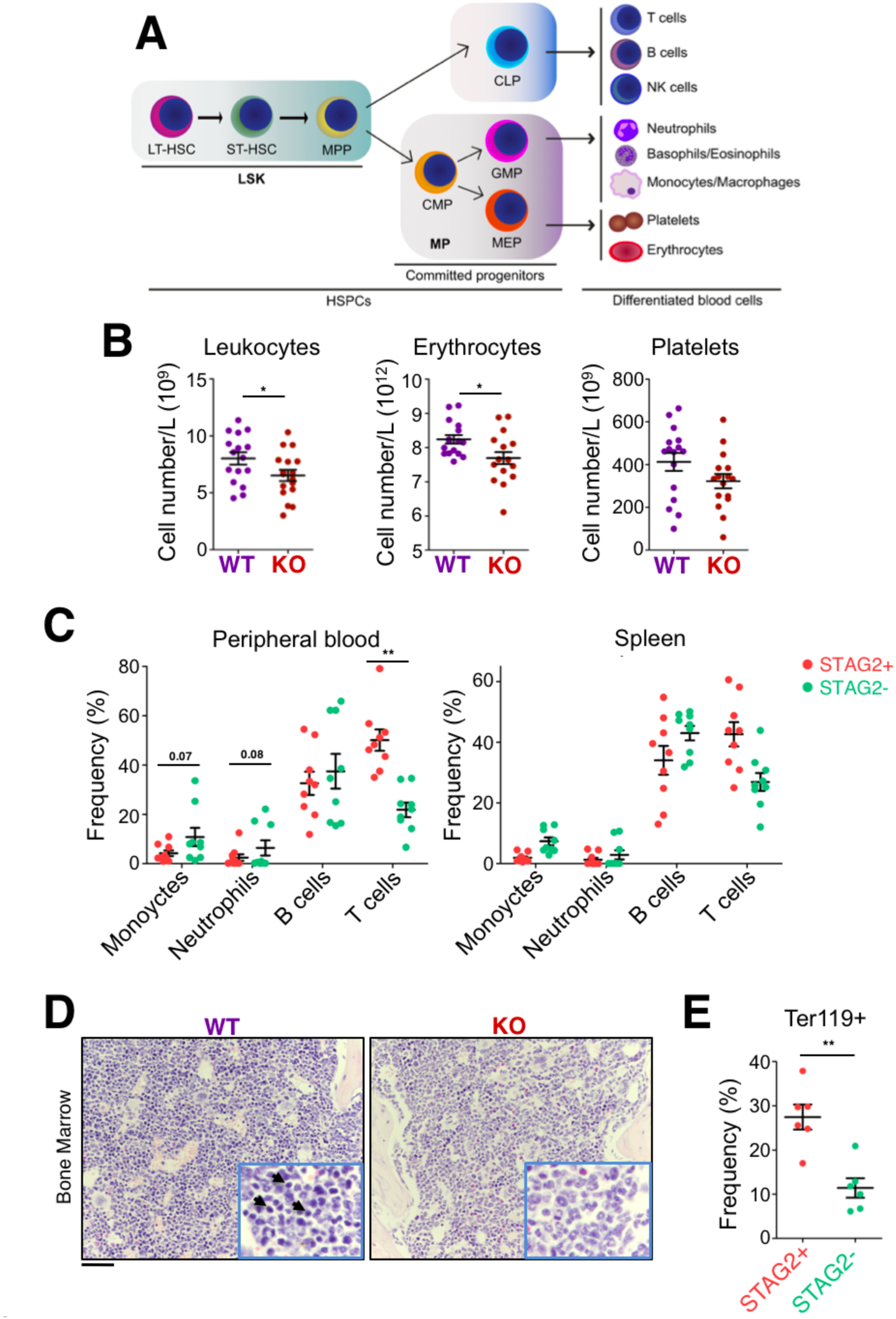
Requirement of STAG2 for normal adult hematopoiesis. **A.** Scheme depicting normal hematopoiesis. **B.** Peripheral blood counts of 12 week-old KO and WT mice (n=16 mice/genotype). Error bars indicate SEM. Two-sided Mann-Whitney U test; *P < 0.05. **C.** Flow cytometry analysis of GFP+ (STAG2-) or Tomato+ (STAG2+) leukocyte populations in peripheral blood and spleen of 12 week-old KO mice (n=9). Monocytes (CD3- B220-CD11b+ Ly6G-); neutrophils (CD3- B220- CD11b+ Ly6G+); B cells (CD3- B220+); T cells (CD3+ B220-). Error bars indicate SEM. Unpaired t test; **P < 0.01. **D.** H-E staining of bone marrow from 12 week-old WT and KO mice shows a decrease in the erythrocyte population (arrows) in KO mice. Scale bar, 50 µm. **E.** Flow cytometry analysis of bone marrow Ter119+ cells (n=6 KO). Error bars indicate SEM. Unpaired t test; **P < 0.01.

**Figure S3.**
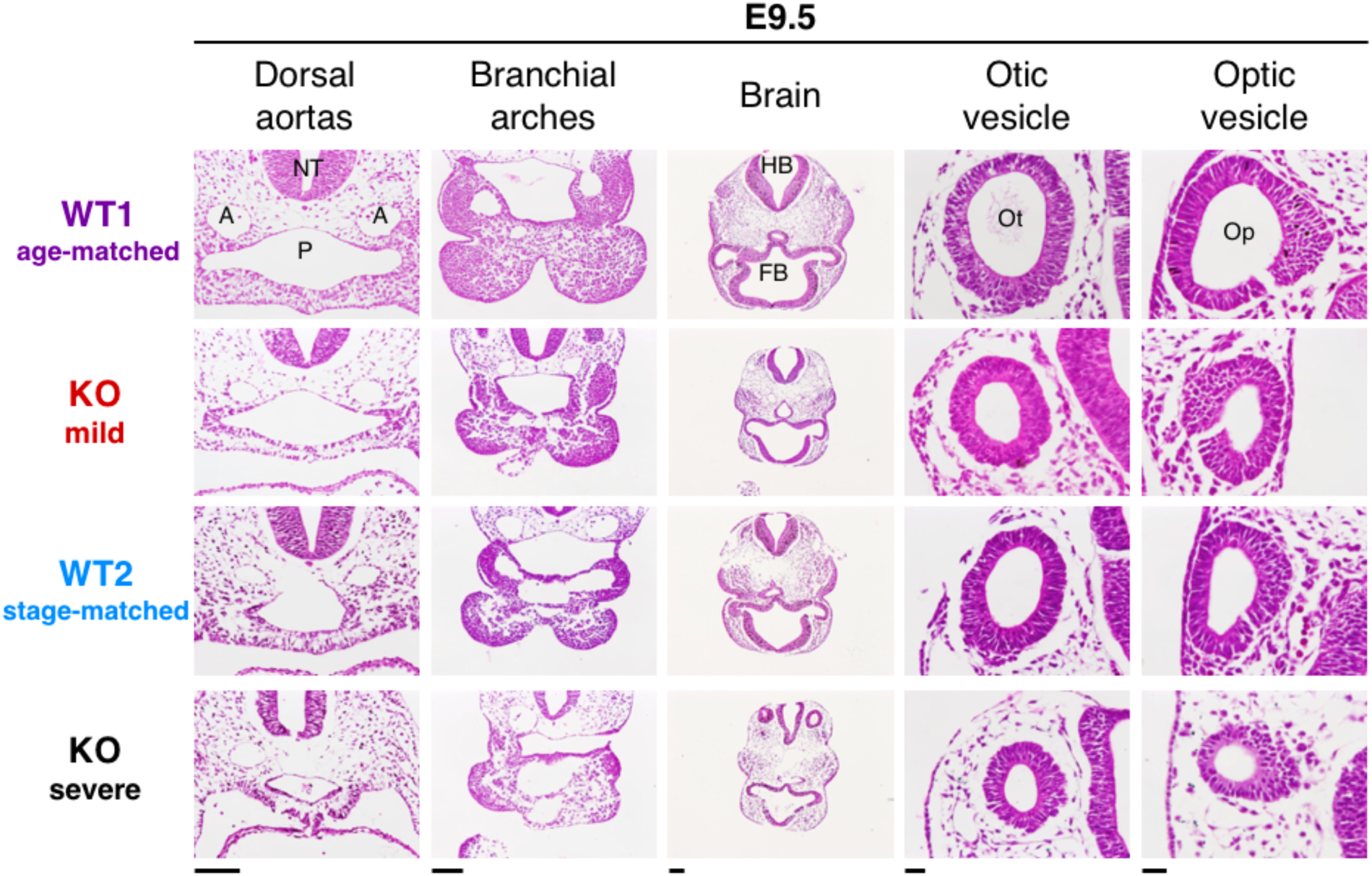
Global developmental defects in *Stag*2 null E9.5 embryos. H-E stained transverse sections of *Stag2* null (KO, mild and severe), WT1 (age-matched control) and WT2 (stage-matched control) embryos extracted at E9.5. NT: neural tube. A: aorta. P: pharynx. HB: hindbrain. FB: forebrain. Ot: otic vesicle. Op: optic vesicle. Scale bars (for entire column): 100 µm for dorsal aortas, branchial arches and brain; 25 µm for otic and optic vesicles.

**Figure S4.**
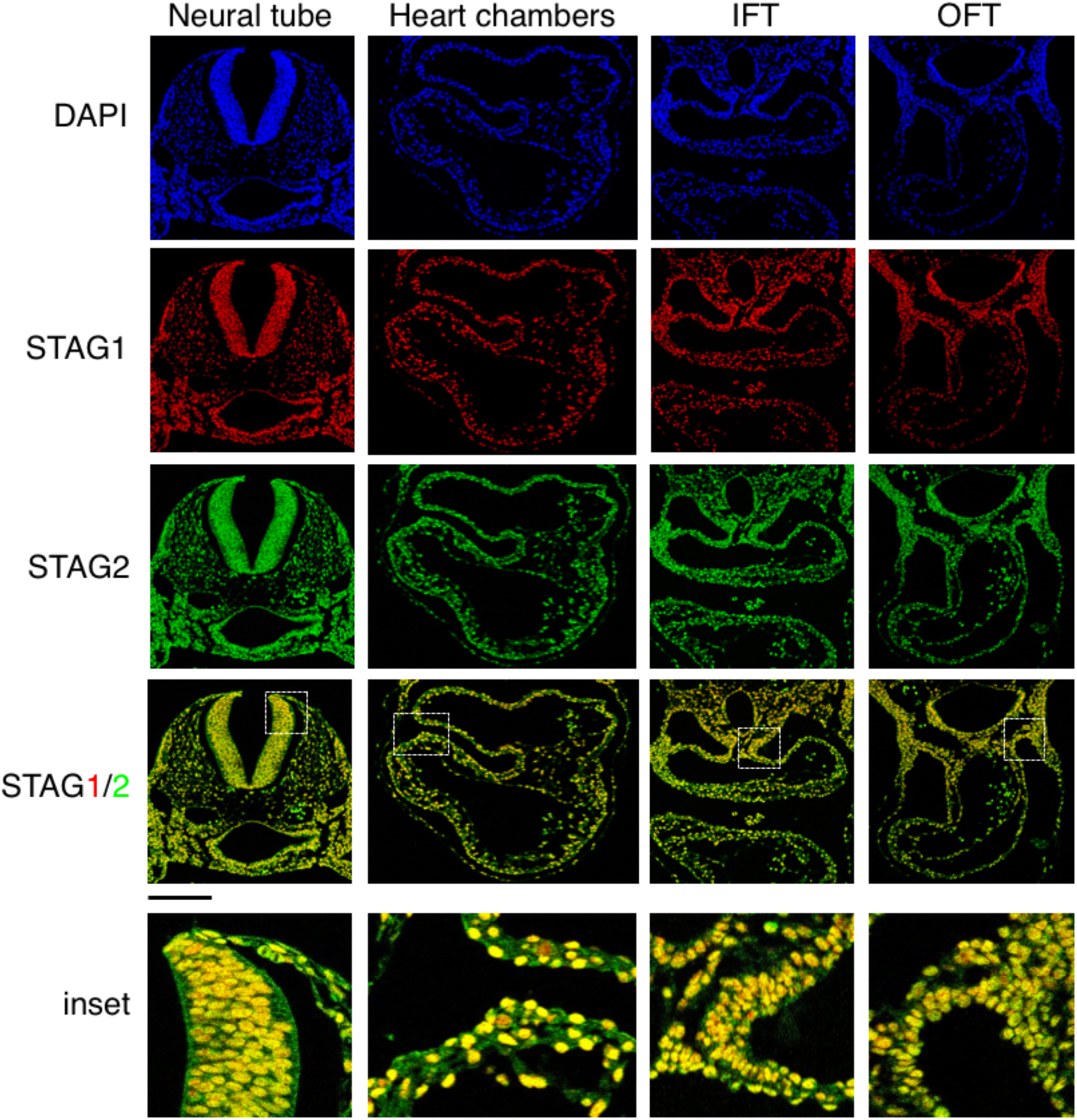
Distribution of cohesin variants in the E9.5 embryo. Immunofluorescence co-staining of STAG1 (red), STAG2 (green) and DAPI (blue) in transverse sections containing the heart of wild type E9.5 embryos. Scale bar, 200 µm.

**Figure S5.**
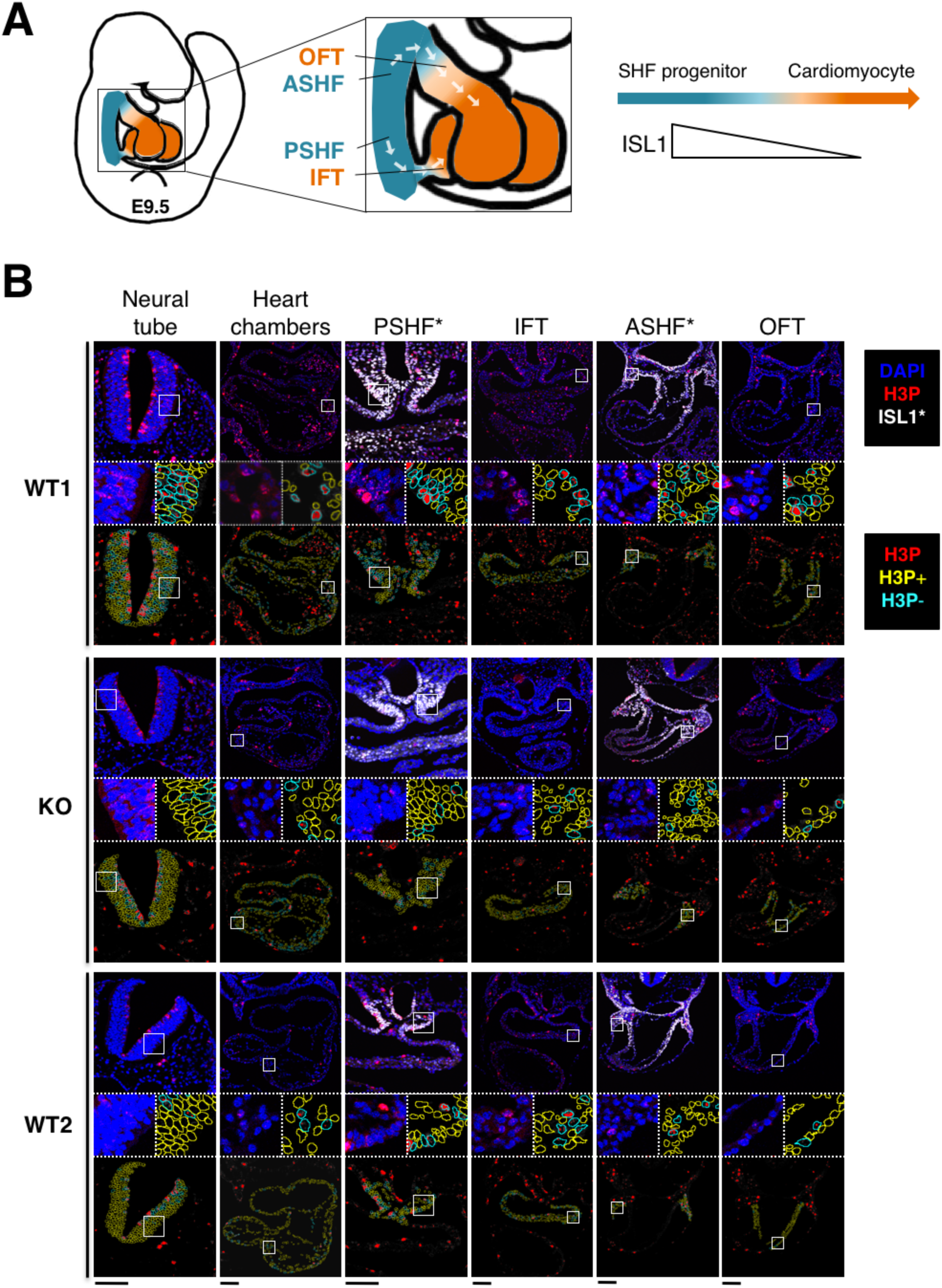
Decreased proliferation in *Stag2* null embryos. **A.** Scheme depicting the migration of second heart field (SHF) progenitors into the heart tube of an E9.5 embryo; outflow tract (OFT), anterior SHF (ASHF), posterior SHF (PSHF), inflow tract (IFT), Islet1 transcription factor (ISL1). **B.** Representative images of H3P staining and image processing in E9.5 WT1 (age-matched control), KO (mild phenotype) and WT2 (stage-matched control) embryos. For each region and genotype are shown the original immunofluorescence signals (top) of DAPI (blue), H3P (red) and ISL1 (white, only shown in regions marked with *), and the binary H3P signal (bottom), with corresponding insets. Scale bars, 100 µm.

**Figure S6.**
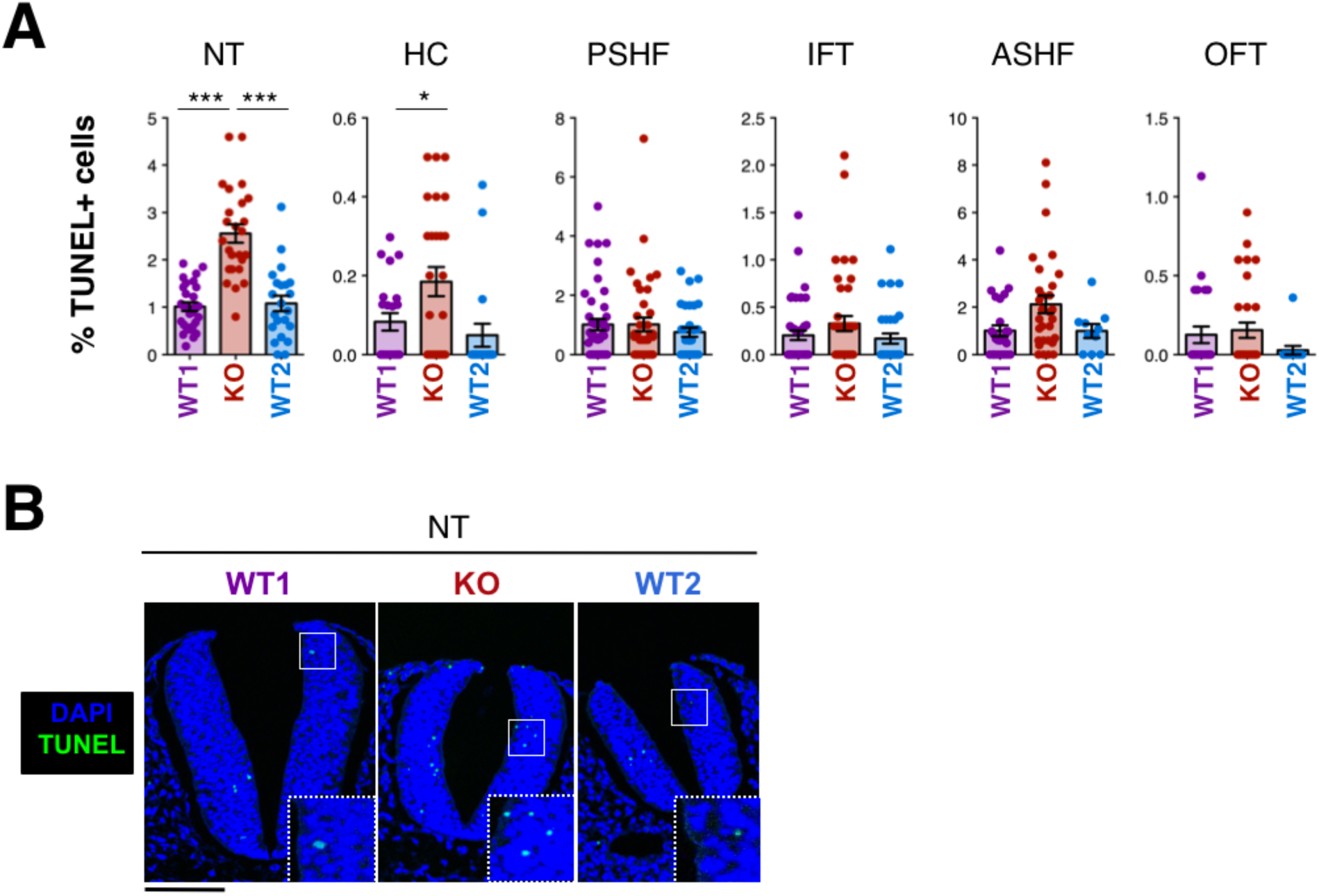
Increased apoptosis in *Stag2* null embryos. **A**. Quantification of TUNEL-positive cells as readout for apoptosis in E9.5 WT1, KO and WT2 embryos in neural tube (NT), heart chambers (HC), posterior secondary heart field (PSHF), inflow tract (IFT), anterior secondary heart field (ASHF) and outflow tract (OFT). At least 10 sections from 3-4 embryos were analyzed per genotype and region. Mean ± SEM are shown. Kruskal-Wallis test and Dunn’s multiple comparison post-test; *** P<0.001, * P<0.05, ns P≥0.05. **B**. Representative images of TUNEL staining in neural tube. Scale bar, 100 µm. Individual dots represent values for one section.

**Fig. S7.**
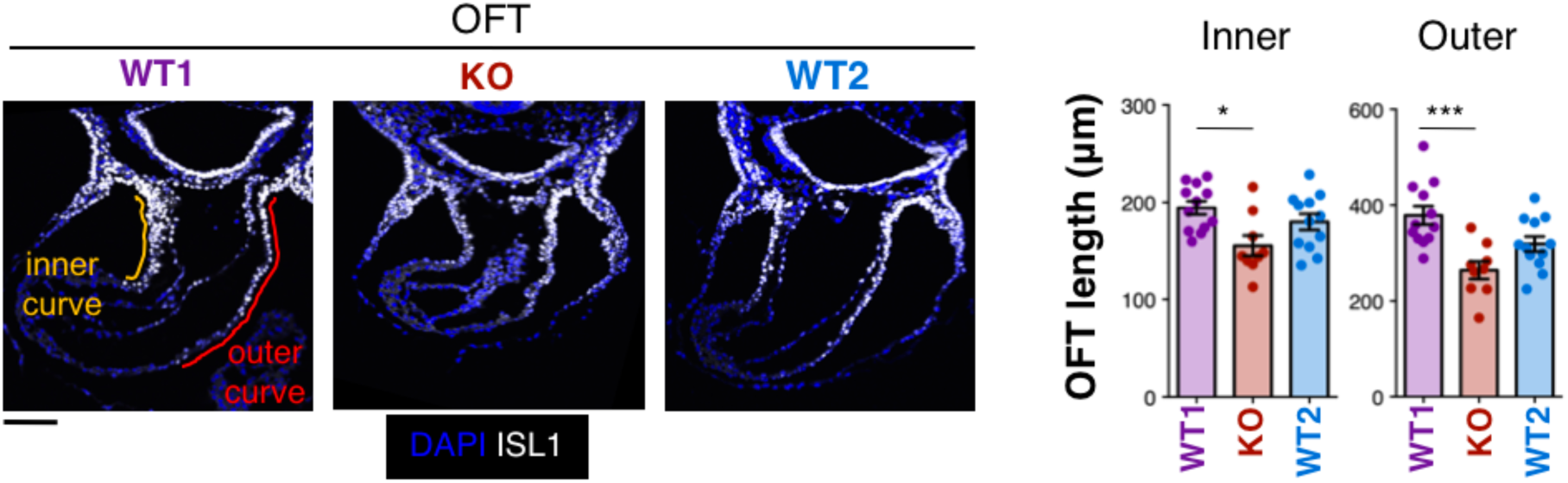
Decreased outflow tract length in *Stag2* null embryos. Representative outflow tract (OFT) images for WT1 (age-matched control), KO (mild phenotype) and WT2 (stage-matched control) E9.5 embryos (left) and measurement of inner and outer OFT length (right). 9-12 sections from 3-4 embryos were analyzed per genotype. Mean ± SEM are shown. Kruskal-Wallis test and Dunn’s multiple comparison post-test; *** P<0.001, * P<0.05, ns P≥0.05.

